# A Human H5N1 Influenza Virus Expressing Bioluminescence for Evaluating Viral Infection and Identifying Therapeutic Interventions

**DOI:** 10.1101/2025.03.28.646035

**Authors:** Ramya S. Barre, Ruby A. Escobedo, Esteban M. Castro, Anastasija Cupic, Mahmoud Bayoumi, Nathaniel Jackson, Chengin Ye, Aitor Nogales, Roy N. Platt, Timothy J. C. Anderson, Adolfo García-Sastre, Ahmed Mostafa, Luis Martinez-Sobrido

## Abstract

A multistate outbreak of highly pathogenic avian influenza virus (HPAIV) H5N1 in dairy cows was first reported on March 25, 2024, in the United States (US), marking the first discovery of HPAIV H5N1 in cattle. Soon after, a dairy worker on an affected dairy farm became the first human case linked directly to this outbreak. Studies with influenza A virus (IAV) require secondary methods to detect the virus in infected cells or animal models of infection. We modified the non-structural (NS) genome segment of the human A/Texas/37/2024 (HPhTX) H5N1 virus to create a recombinant virus expressing nanoluciferase (HPhTX NSs-Nluc), enabling the tracking of virus in cultured cells and mice via *in vitro*, *ex vivo*, and *in vivo* imaging systems (IVIS). *In vitro*, HPhTX NSs-Nluc showed growth and plaque characteristics similar to its wild-type (WT) counterpart. *In vivo*, HPhTX NSs-Nluc allowed tracking viral infection in the entire animals and in the organs of infected animals using *in vivo and ex vivo* IVIS, respectively. Importantly, the morbidity, mortality, and replication titers of HPhTX NSs-Nluc were comparable to those of the WT HPhTX. *In vitro*, HPhTX NSs-Nluc was inhibited by Baloxavir acid (BXA) to levels observed with WT HPhTX. We also demonstrate the feasibility of using HPhTX NSs-Nluc to evaluate the antiviral activity of BXA *in vivo*. Our findings support that HPhTX NSs-Nluc represents an excellent tool for tracking viral infections, including the identification of prophylactics or therapeutics for the treatment of the HPAIV H5N1 responsible of the outbreak in dairy cows.

## INTRODUCTION

Influenza viruses are negative-sense, single-stranded RNA viruses belonging to the family *Orthomyxoviridae* (1). Influenza viruses are classified in four types: A, B, C, and D (IAV, IBV, ICV, and IDV, respectively). IAVs have been responsible for seasonal epidemics and occasional pandemics of significant public health concern in humans (1). The highly pathogenic avian influenza virus (HPAIV) H5N1 of clade 2.3.4.4b has been responsible for outbreaks in wild and domestic birds (2). Recent reports have shown the wide distribution and transmissibility of an emerging HPAIV H5N1 clade 2.3.4.4b, genotype B3.13, in cattle and humans in the United States (US), representing a threat for avian and other mammalian species, including cats, raccoons, opossums, dairy cattle, and humans (3). In cattle, virus replication mainly takes place in the mammary glands (3, 4). Between March 2024 and March 2025, a total of 70 HPAIV H5N1 human cases, including one death, have been reported in the US (5). Despite that the potential public risk of transmission and adaptation in humans of HPAIV H5N1 viruses is considered low, there is a risk of human infections among poultry and dairy farm workers, and veterinarians. The virus has also caused damage to the poultry and dairy industry, affecting the food production chain. The ability of HPAIV to infect and transmit in cows is not restricted to HPAIV H5N1 clade 2.3.4.4b genotype B3.13 as evidenced by more recent independent introductions of another genotype, D1.1, in dairy cows in US (6).

Studying IAV involves assessing the presence of the virus in infected cells or animal models of infection. Current methods to detect IAV *in vitro, ex vivo,* and *in vivo* are often laborious and time-consuming. Many studies, including ours (7–10), have shown that recombinant viruses, including IAV, expressing reporter genes represent an efficient approach for assessing the presence of viruses in cultured cells and in animal models. We have previously described IAV expressing fluorescent proteins from the NS segment (9, 11, 12). Nanoluciferase (Nluc) is a bioluminescent protein that allows tracking viruses using *in vivo* imaging systems (IVIS), while fluorescent proteins can usually only be assessed by *ex vivo* imaging, as the background limits the sensitivity of detection using IVIS. We have previously demonstrated the feasibility of using recombinant viruses expressing Nluc for tracking viral infections in different animal models, including mice and hamsters, and their feasibility to facilitate the identification of prophylactics and therapeutics for the treatment of viral infections (7–9, 11, 13).

In this study, a HPAIV A/Texas/37/2024 H5N1 (HPhTX) was engineered to express Nluc from the NS segment (HPhTX NSs-Nluc) by fusing to the C-terminal of the viral NS1. Since NS1 is one of the most abundant proteins expressed during IAV infection, fusion of Nluc facilitates efficient expression of the bioluminescent reporter *in vitro, ex vivo,* and *in vivo*. *In vitro*, HPhTX NSs-Nluc exhibited plaque phenotypes, replication kinetics, and susceptibility to anti-influenza drugs comparable to those of the wild-type (WT) HPhTX. *In vivo,* HPhTX NSs-Nluc mirrored pathogenicity and ability to replicate similar to WT HPhTX. Notably, we were able to monitor HPhTX NSs-Nluc in the entire mice using IVIS and demonstrate the feasibility of HPhTX NSs-Nluc to identify therapeutics. In addition, Nluc expression was detectable *ex vivo* in the nasal turbinate (NT), lungs, and brains of infected mice. Importantly, Nluc expression remained stable for 10 consecutive passages *in vitro.* Altogether, HPhTX NSs-Nluc provides a valuable platform for investigating HPhTX *in vitro* and *in vivo*, enabling real-time viral tracking via live imaging systems, and including the efficient detection of prophylactics or therapeutics for the prevention or treatment of HPhTX infection in cell culture and validated animal models of infection.

## MATERIALS AND METHODS

### Biosafety

All *in vitro* and *in vivo* studies involving HPAIV A/Texas/37/2024 H5N1 were carried out in BSL-3 and ABSL-3 laboratories at Texas Biomedical Research Institute (Texas Biomed). Studies were approved by both the Institutional Biosafety Committee (IBC) and the Institutional Animal Care and Use Committee (IACUC) at Texas Biomed.

### Cells and Viruses

MDCK (Madin-Darby canine kidney) and human 293T cells were grown in Dulbecco’s modified Eagle’s medium (DMEM, Gibco) supplemented with 10% fetal bovine serum and 1% PSG (penicillin,100 units/ml; streptomycin 100 μg/ml; L-Glutamine, 2 mM) at 37°C in a 5% CO_2_ cell culture incubator. HPhTX, HPhTX NSs, and HPhTX NSs-Nluc were grown in MDCK cells as previously described (14). Viral titers were determined by standard plaque assay in MDCK cells.

### Plasmids and viral rescues

Recombinant A/Texas/37/2024 H5N1 (HPhTX), where Nluc was fused to the C-terminus of NS1 was engineered and generated as previously described (15). The modified NS segment was synthesized *de novo* (Bio Basic, USA) with the appropriate restriction sites for subcloning into the ambisense plasmid pHW2000 to generate the pHW-H5N1-NSs_AgeI/NheI plasmid. The recombinant NSs segment contains the NS1 open reading frame (ORF) without stop codons or splice acceptor sites, followed by AgeI and NheI restriction sites, the porcine teschovirus-1 (PTV-1) 2A autoproteolytic cleavage site (ATNFSLLKQAGDVEENPGP) and the entire ORF of the nuclear export protein, NEP (13). The Nluc ORF was cloned using the AgeI and NheI sites into pHW-H5N1-NSs_AgeI/NheI to generate the pHW-H5N1-NSs_Nluc plasmid for virus rescue. Plasmid constructs were confirmed by DNA sequencing (Plasmidsaurus). Recombinant HPhTX, HPhTX NSs, and HPhTX NSs-Nluc viruses were rescued as previously described (14, 16–19). A recombinant low pathogenic A/Texas/37/2024 H5N1 lacking the NS1 protein (LPhTXdNS1) containing a mutated HA protein with a monobasic cleavage site (PQIETR/GLF) was generated and propagated as described previously (11). Rescued viruses were plaque purified and amplified in MDCK cells to generate viral stocks. Viral stocks were aliquoted and stored at −80°C until further use.

### Cell-based interferon (IFN) bioassay

To evaluate the activation of IFNβ promoter *in vitro*, MDCK cells expressing the green fluorescent protein (GFP) and firefly luciferase (FFluc) protein under the control of the IFNβ promoter (MDCK pIFNβ-GFP/IFNβ-FFluc) (7, 20) were cultured in 12-well plates (5 x 10^5^ cells/well, triplicates) and mock-infected or infected (multiplicity of infection, MOI 1) with HPhTX, HPhTX NSs, and HPhTX NSs-Nluc for 12 h. The LPhTXdNS1 was used as a control. IFNβ promoter activation was evaluated by GFP expression using fluorescent microscope and by FFluc activity from cell lysates using Promega luciferase reporter assay and a Glowmax microplate reader. Nluc expression in cells infected with HPhTX NSs-Nluc was determined by assessing Nluc expression in the cell culture supernatants using the Promega Nluc assay kit and a Glowmax microplate reader. Viral infections were confirmed by immunostaining the cells with a monoclonal antibody (HT103) against the viral nucleoprotein (NP) (21) and an Alexa Fluor® 594 AffiniPure™ goat anti-mouse IgG (H+L) secondary antibody.

### Virus growth kinetics

Multicycle growth kinetics were conducted in MDCK cells (6-well plate format, 10^6^ cells/well, triplicates). Cell monolayers were infected (MOI of 0.00001) with the indicated viruses. After 1h of viral adsorption at room temperature (RT), cells were overlayed with DMEM 0.3% BSA, 1% PSG and were incubated at 37°C in a 5% CO_2_ incubator. At 12, 24, 48, and 72 h post-infection (hpi), viral titers in the cell culture supernatants were determined by plaque assay as previously described (14). The presence of Nluc in the same cell culture supernatants was quantified using the Promega Nluc assay kit and a Glowmax microplate reader. Mean values and standard deviation (SD) were calculated using Microsoft Excel Software.

### Plaque assay and immunostaining

Confluent monolayers of MDCK cells (6-well plate format, 10^6^ cells/well) were infected with HPhTX, HPhTX NSs, or HPhTX NSs-Nluc viruses. After 1 h viral adsorption, infectious virus was replaced by infection media containing agar and the plates were incubated at 37°C in a 5% CO_2_ incubator. At 48 and 72 h, plates were fixed overnight in 4% formaldehyde. For visualization of Nluc expression, the agar overlay was removed, and the plates were incubated with Nluc substrate in 1X PBS and imaged using a Chemidoc. Following Nluc visualization, plates were permeabilized with 0.5% Triton X-100 in 1X PBS for 15 min at RT and immunostained with the NP MAb HT103 (21). Immunostaining was developed using a Vectastain ABC kit and a DAB HRP substrate kit, following manufacturer’s recommendations.

### Genetic stability of HPhTX NSs-Nluc *in vitro*

MDCK cells (6-well plate format, 10^6^ cells/well) were infected (MOI of 0.01) with HPhTX NSs-Nluc and incubated at 37°C in a 5% CO_2_ incubator until ∼70-80% cytopathic effect (CPE) was observed. Cell culture supernatants were then harvested and diluted (1:100) for subsequent serial passages in fresh MDCK cells (6-well plate format, 10^6^ cells/well) for a total of 10 passages. At each passage, cell culture supernatants were used to assess Nluc expression and viral titration using plaque infectivity assay. Viral RNA from passage 1, 5 and 10 were extracted using TRIzol^TM^ reagent (Thermo fisher Scientific, US), for whole genome sequencing using next-generation sequencing platform, MinION (Oxford Nanopore Technologies). Sample libraries were prepared using the Native Barcoding Kit 24 V14 (SQK-NBD114.24, Oxford Nanopore Technologies) and ran on R10.4.1. Flow Cells (FLO-MIN114, Oxford Nanopore Technologies) per manufacturer’s instructions. The read length stats with n50 v1.7.0 then the raw reads were trimmed using nanoq v0.10.0 (22). Bases with quality PHRED quality scores less than 7 were removed along with the first and last 25bp of each read. Reads less than 500 bp after trimming were removed from downstream analyses. Filtered reads were mapped to the HPhTX NSs-Nluc reference sequence using minimap2 v2.28-r1209 (23) and the ‘-x map-ont’ option for mapping long error prone nanopore reads. Mapping rates were calculated with SAMtools flagstat v1.21 (24). We estimated coverage across each genome with MosDepth v0.3.10 (25). We re-assessed indel quality scores and called variants using LoFreq v2.1.5 (26). We limited depth of coverage (--max-depth) for variant calling at 10,000x and removed the default filters applied by LoFreq. Rather, we removed any variants that were present in less than 25% of reads or regions of the genome that contained less than 100x read depth.

### Antiviral assays

Antiviral assays were conducted as previously described (27). Briefly, confluent monolayers of MDCK cells (96-well plate format, 5×10^4^ cells/well, quadruplicates) were infected with 100 plaque-forming units (PFU) of HPhTX NSs-Nluc and incubated for 1 h at 37°C. After viral adsorption, two-fold serial dilutions of BXA (starting concentration of 10 mM) in infection media (DMEM 0.3% BSA, 1% PSG) were added to each well and plates were incubated at 37°C in a 5% CO_2_ incubator. At 48 hpi, the supernatants were collected for Nluc activity measurements, and the cell monolayers were fixed with 10% paraformaldehyde (PFA) and stained at RT with 1% crystal violet for 20 min. Once dried, stained monolayers with crystal violet were destained with 200µl absolute methanol for 10 min and measured at λmax 570 nm as described previously (14). Cystal violet and Nluc values in virus-infected cells in the absence of BXA were used to calculate 100% viral infection. Mock-infected cells were used as negative controls to calculate Nluc background levels and 100% viral inhibition as measured by crystal violet values. The average of the quadruplicate wells was used to calculate the SD of neutralization using Microsoft Excel Software. The 50% inhibition concentration (IC_50_) was calculated using a sigmoidal dose-response curve (GraphPad Prism software).

### Mouse experiments

Six-week-old female C57BL/6 mice were purchased from The Jackson Laboratory and housed under specific pathogen-free conditions at Texas Biomed. For viral infection, mice were anesthetized intraperitoneally (i.p.) with a mixture of Ketamine (100 mg/mL) and Xylazine (20 mg/mL). Mice were intranasally (i.n.) inoculated with 50 µL of either HPhTX NSs or HPhTX NSs-Nluc, with doses of 10, 10², 10³, and 10 PFU per mouse. *In vivo* Nluc imaging was performed using the IVIS Spectrum multispectral imaging system. Mice were anesthetized on days 1, 2, 4, 6, and 8 and injected retro-orbitally with 100 µL of Nano-Glo luciferase substrate (Promega, US) diluted to 1:10 in 1X PBS. Immediately following injection, bioluminescence was measured using IVIS. Bioluminescence data were acquired and analyzed using the Aura program (AMI Spectrum). Morbidity, as indicated by changes in body weight, and mortality, as assessed by survival rate, were monitored over 14 days. Mice that lost 25% or more of their initial body weight were considered to have reached the experimental endpoint and were humanely euthanized. Survival data were analyzed using Kaplan-Meier curves. The 50% mouse lethal dose (MLD_50_) was calculated by assessing the survival rates at doses ranging from 10 to 10 PFU/ml. For *ex vivo* imaging, lungs and brains were collected following euthanasia. After imaging, lungs and brains, and NT collected at necropsy, were homogenized in 1 mL of 1X PBS using a Precellys tissue homogenizer (Bertin Instruments) for 20 seconds at 7,000 rpm. Tissue homogenates were then centrifuged at 12,000 x g at 4°C for 5 min, and the supernatants were collected for titration by plaque assay, as described above. Supernatants collected from tissue homogenates were also used to assess levels of Nluc using the Nano-Glo luciferase substrate and a Glowmax microplate reader.

Likely, six-week-old female C57BL/6 mice (n=5) were purchased from The Jackson Laboratory and housed under specific pathogen-free conditions at Texas Biomed. The mice were treated with 15 mg/kg baloxavir marboxil at 6 h before intranasal challenge infection with 10^2^ PFU of HPhTX NSs-Nluc virus. The mice were monitored for bodyweight and survival rates for 14 days post-infection and subjected to IVIS at days 2, 4, 6, and 8 post-infections. To investigate the impact of treatment on viral replication, necropsy groups of mice (n=4) were subjected to IVIS at day 6 post-infection and necropsied to collect the nasal turbinates, lung and brain tissues for *ex vivo* imaging and viral load assessment using plaque assay and Nluc expression levels.

### Statistical analyses

All the graphs, calculations, and statistical analyses were performed using GraphPad Prism software version 9.5.1 (GraphPad Software, LLC, USA).

## RESULTS

### Generation and *in vitro* characterization of HPhTX NSs-Nluc

To create a replication-competent HPhTX expressing Nluc (HPhTX NSs-Nluc), the NS segment was modified to encode non-overlapping NS1 and NEP ORFs as previously described (9, 11). NS1 and NEP ORFs were separated by the porcine teschovirus-1 (PTV-1) 2A autoproteolytic cleavage site, enabling the independent expression of both ORFs (**Fig 1A**). A recombinant HPhTX virus expressing a split NS segment (NSs) without the fusion of Nluc to the C-terminal of NS1, referred to as HPhTX NSs, was also generated and used as control in the experiments (**Fig 1A**). AgeI and NheI restriction sites were incorporated after the 2A site to insert the Nluc ORF fused the C-terminal endo of NS1 to generate the Nluc-expressing recombinant HPhTX, designated HPhTX NSs-Nluc (**Fig 1B**). After viral rescue, fitness of HPhTX, HPhTX NSs, and HPhTX NSs-Nluc were assessed in monolayers of infected (MOI of 0.00001) MDCK cells at different times post-infection (**Fig 1C**). Nluc expression was also measured from the same cell culture supernatants, which was detectable only in the supernatants of HPhTX NSs-Nluc-infected MDCK cells (**Fig 1C**). We also compared the plaque phenotype of HPhTX, HPhTX NSs, and HPhTX NSs-Nluc in MDCK cells at 48 and 72 hpi and demonstrate Nluc expression in the cells infected with HPhTX NSs-Nluc (**Fig 1D**). The plaque phenotypes of the three viruses were comparable at 48 and 72 hpi, which aligned with the similar replication kinetics at the same times after infection. Altogether, these results demonstrate the feasibility of generating replicating competent HPhTX expressing a modified NSs segment where the NS1 and NEP ORFs are not overlapping and the feasibility of using this modified NSs strategy for the expression of Nluc fused to the NS1 without significantly affecting the growth kinetics or plaque phenotype in MDCK cells.

**Figure 1.**
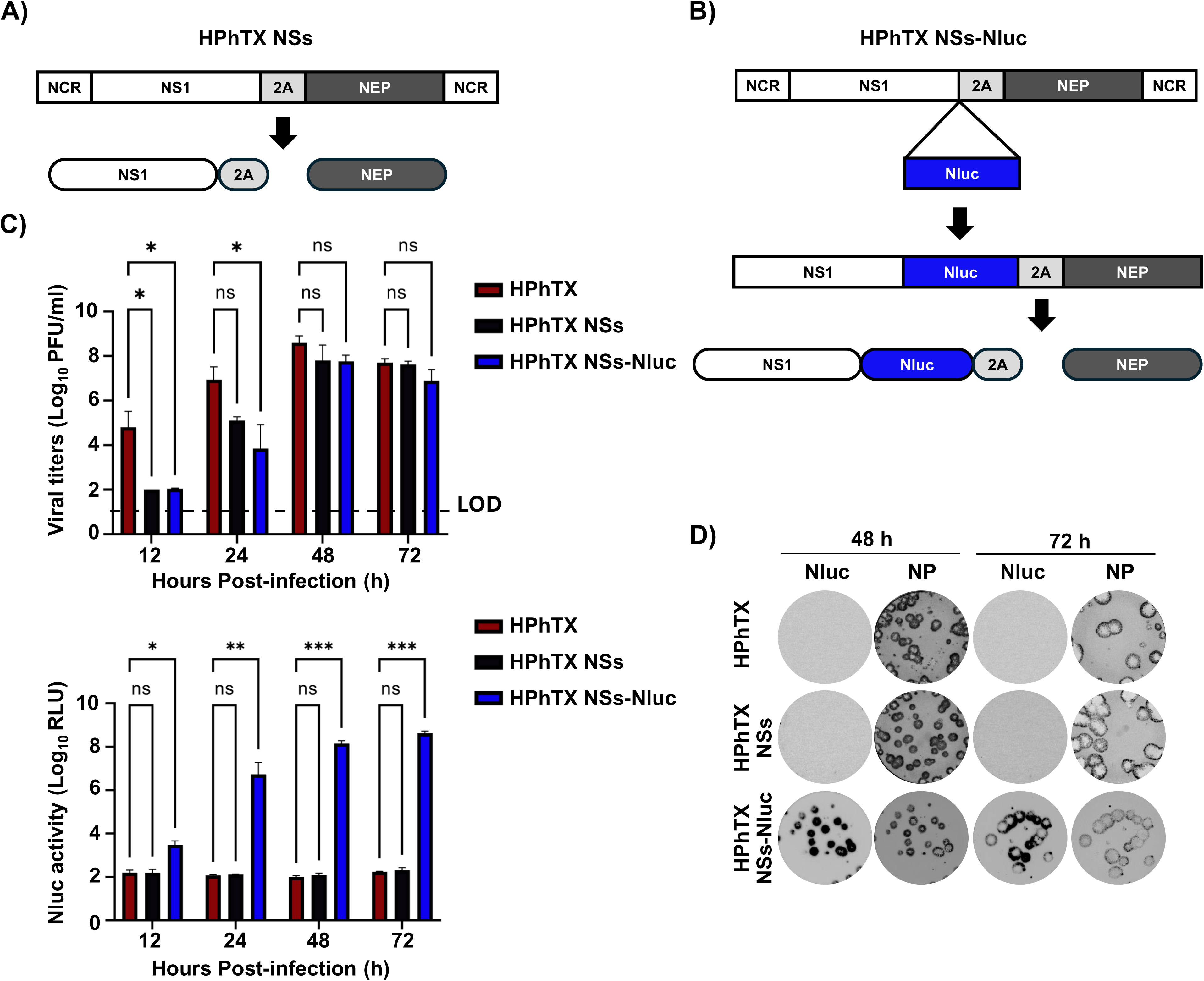
*In vitro* characterization of HPhTX NSs-Nluc. (**A-B**) Schematic representation of HPhTX NSs (**A**) and HPhTX NSs-Nluc (**B**) viral segments. The viral NS1 and NEP ORFs, and PTV-1 2A sequences are represented in white, black, and gray, respectively. The Nluc ORF is represented in blue. (**C**) Multicycle growth kinetics of HPhTX, HPhTX NSs, and HPhTX NSs-Nluc as determined by plaque assay in MDCK cells (top); and Nluc activity from same cell culture supernatants (bottom). Data represents means and SD for triplicates. (**D**) Plaque phenotype of HPhTX, HPhTX NSs, and HPhTX NSs-Nluc in MDCK cells. Viral plaques were evaluated at 48 and 72 hpi via staining with Nluc substrate (left) and NP immunostaining (right). A two-way repeated measure ANOVA with Geisser-Greenhouse correction. Post-hoc multiple comparisons performed using Šídák or Bonferroni method to compare groups within each time-point. The significant differences are indicated (ns=non-significant, * = *p* < 0.05, ** = *p* < 0.01, *** = *p* < 0.001, **** = *p* < 0.0001).

### HPhTX NSs-Nluc efficiently inhibits IFN**β** promoter activation

During viral infection, the IAV NS1 protein is involved in inhibiting IFN and host antiviral responses (28). To demonstrate that the modified NSs segment in HPhTX NSs, or the fusion of Nluc to the C-terminal of NS1 protein in HPhTX NSs-Nluc did not impact its ability to control IFN responses, MDCK IFNβ-GFP/IFNβ-FFluc cells, which express GFP and FFluc reporter genes under the control of the IFNβ promoter, were infected (MOI of 1) with HPhTX, HPhTX NSs, or HPhTX NSs-Nluc. As an internal control, MDCK IFNβ-GFP/IFNβ-FFluc cells were infected at the same MOI with a LPhTX lacking NS1 (LPhTXdNS1), or mock-infected. After 12 hpi, we evaluated IFNβ promoter activation by analyzing GFP and FFluc expression using fluorescent microscopy and a luciferase plate reader, respectively. Expression of GFP and FFluc was readily detected in MDCK IFNβ-GFP/IFNβ-FFluc cells infected with LPhTXdNS1 (**Figs. 2A and 2B**). However, we did not observe activation of the IFNβ promoter in MDCK IFNβ-GFP/IFNβ-FFluc cells infected with HPhTX, HPhTX NSs, or HPhTX NSs-Nluc with comparable levels of infection, as determined by NP staining (**Figs. 2A and 2B**). These results suggests that HPhTX NSs and HPhTX NSs-Nluc efficiently inhibit IFNβ promoter activation, similar to HPhTX. We also determined Nluc expression in cell culture supernatants from MDCK IFNβ-GFP/IFNβ-FFluc infected cells (**Fig. 2C**). As expected, Nluc expression was only observed in the supernatant of cells infected with HPhTX NSs-Nluc (**Fig. 2C**).

**Figure 2.**
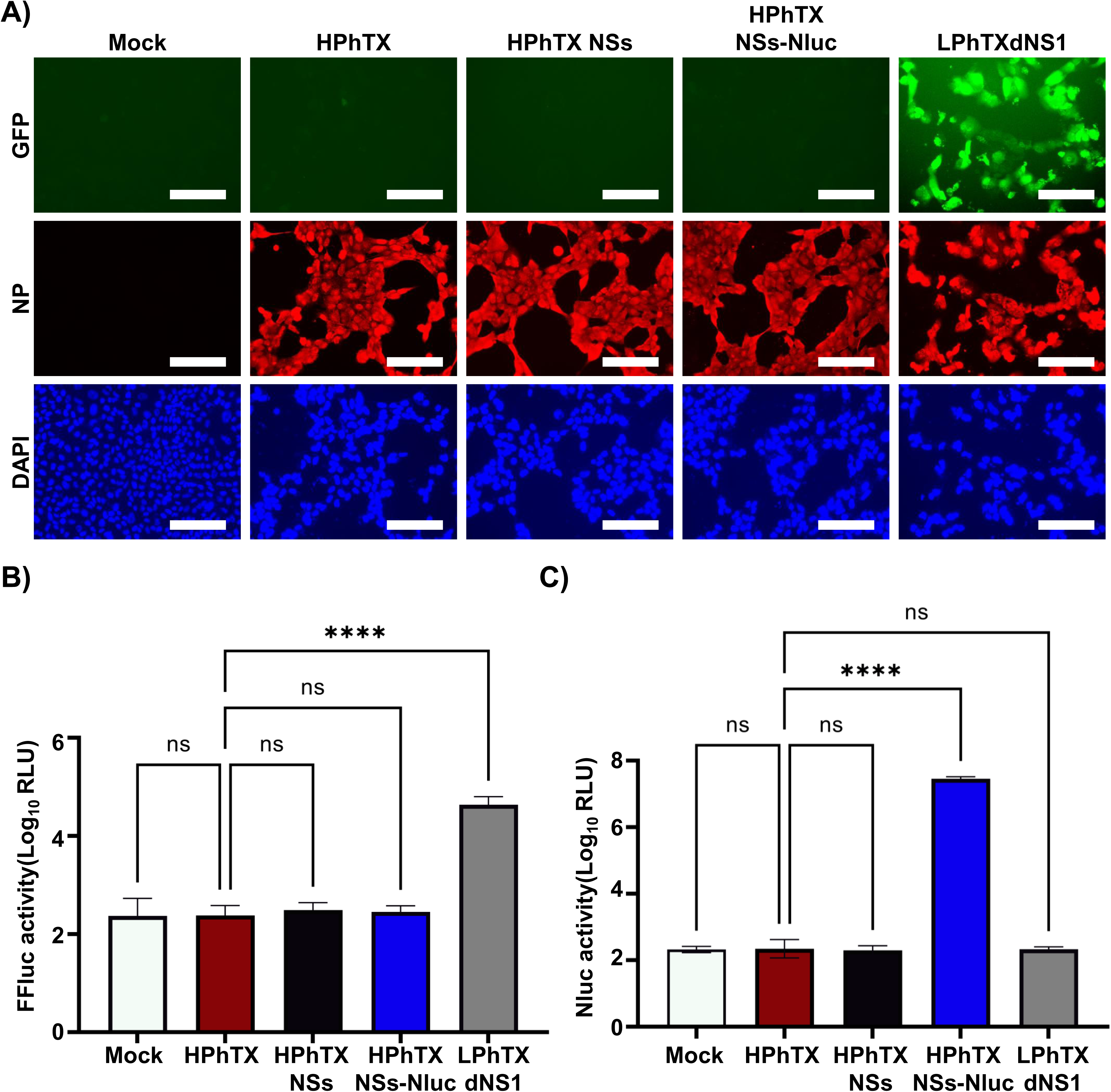
HPhTX NSs-Nluc inhibits ifNβ promoter activation. MDCK IFNβ-GFP/IFNβ-FFluc cells were mock-infected or infected (MOI 1) with HPhTX, HPhTX NSs, HPhTX NSs-Nluc, or LPhTXdNS1 (control). At 12 hpi, IFNβ promoter activation was analyzed by GFP (**A**) and FFluc (**B**) expression. Nluc expression was evaluated in the cell culture supernatants of infected cells (**C**). A Welch’s one-way ANOVA with Geisser-Greenhouse correction. Post-hoc multiple comparisons performed using Dunnett method to compare groups within each time-point. The significant differences are indicated (ns=non-significant, **** = *p* < 0.0001).

### Phenotypic and genetic stability of HPhTX NSs-Nluc *in vitro*

To evaluate the phenotypic and genetic stability of HPhTX NSs-Nluc *in vitro*, the virus was subjected to 10 serial passages in MDCK cells. Nluc expression levels were measured across all passages, with the results demonstrating consistent and stable Nluc expression (**Fig 3A**). The plaque phenotype and Nluc expression was also assessed from viruses collected from cell culture supernatants from passages 1, 5, and 10 (**Fig 3B**). We observed a comparable plaque phenotype across passages 1, 5, and 10 (**Fig 3B**). Importantly, we observed a correlation between the number of Nluc positive plaques and viral NP positive plaques, demonstrating that up to 10 serial passages in MDCK cells, HPhTX NSs-Nluc retained Nluc expression (**Fig 3B**). Genetic stability of HPhTX NSs-Nluc at passages 1, 5, and 10 was confirmed via nanopore sequencing (**Fig 3C**). The summary statistics for the Nanopore long read data are provided in **Table 1**. The NSs-Nluc gene, specifically, had more than 100x coverage across its entire length in all three samples (**Fig 3C**). We identified 1 variant in the NSs-Nluc gene after removing low frequency allele (<25%) (**Table 2**; **Fig 3D**). The G to A mutation at position 1,538 in NEP was present at all passages, but the frequency decreased in the latest passage (**Fig 3D**).

**Figure 3.**
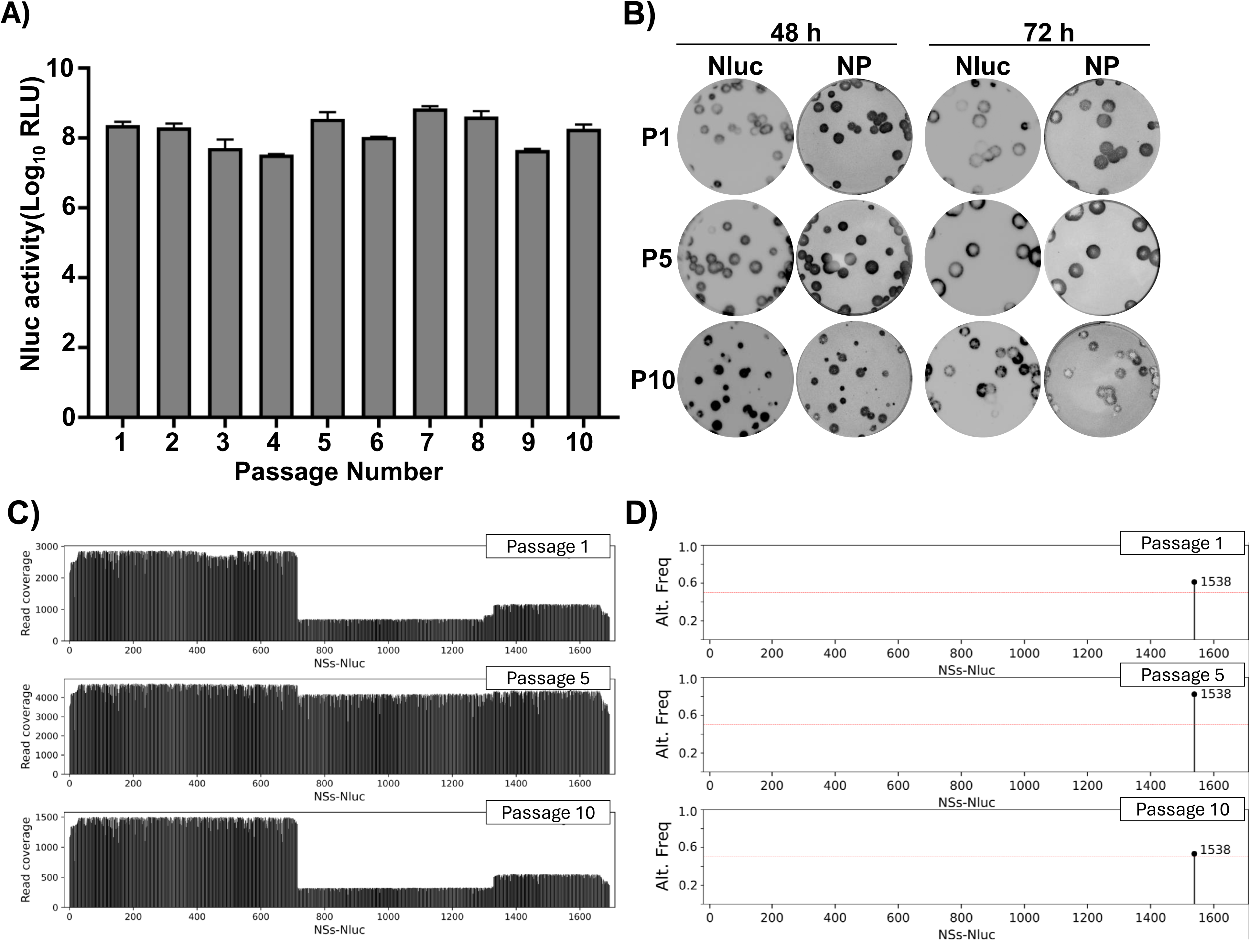
Stability of HPhTX NSs-Nluc *in vitro*. MDCK cells were infected with serial passages of HPhTX NSs-Nluc. (**A**) Nluc expression in cell culture supernatants was evaluated for each of the viral passages. (**B**) Plaque assays of HPhTX NSs-Nluc collected at passages 1, 5, and 10 stained with Nluc substrate (left) or immunostained with the NP MAb HT103 (right). (**C**) NGS sequencing data covering the NSs-Nluc segment across passages 1, 5 and 10. (**D**) Non-reference allele frequency – The passaged samples were compared to the NSs-Nluc reference sequence to identify variants. A single nucleotide variant (SNV) allele frequency from passage 1, 5, and 10 are shown with circles. One variant HPhTX NSs-Nluc (nucleotide 1,538) was at high frequency in all three samples. Variants <25% frequency are not shown. Allele frequencies are provided in Table 2. The red line indicates 25% allele frequency.

**Table 1.** Data description – Sample summary and genome coverage.

| Passage | n Reads | n50 | Gb | Alignment Rate | Mean Coverage | % <100x |
| --- | --- | --- | --- | --- | --- | --- |
| 1 | 98974 | 1,110 | 91.03 | 99.32% | 4,868 | 89.35 |
| 5 | 136007 | 1,159 | 141.98 | 99.89% | 7,978 | 84.7 |
| 10 | 29084 | 1,154 | 30.76 | 98.60% | 1,750 | 77.26 |

**Table 2.** Variant frequencies – All variants at <25% allele frequency are shown on the NSs-Nluc segment.

| Passage | Segment | Pos. | Ref. allele<br>(amino acid) | Alt. allele<br>(amino acid) | Alt. freq | Read depth |
| --- | --- | --- | --- | --- | --- | --- |
| 1 | NSs-Nluc | 1538 | G (Leucine) | A (Leucine) | 61.11% | 1,170 |
| 5 | NSs-Nluc | 1538 | G (Leucine) | A (Leucine) | 82.18% | 4,371 |
| 10 | NSs-Nluc | 1538 | G (Leucine) | A (Leucine) | 53.26% | 552 |

The G to A mutation at position 1,538 is silent and does not result in a change to the corresponding amino acid (leucine). Overall, Nluc sequence results from HPhTX NSs-Nluc passages remained intact without deletions and/or insertions. These findings confirm the stability of HPhTX NSs-Nluc up to 10 passages in MDCK cells.

### HPhTX NSs-Nluc infection in C57BL/6J mice

The pathogenicity of HPhTX NSs-Nluc was assessed in C57BL/6J mice and compared to HPhTX NSs virus (**Fig. 4**). We have previously demonstrated that HPhTX is highly pathogenic in mice with an estimated MLD_50_ of < 10 PFU (14). To that end, mice were i.n. inoculated with 10, 10^2^, 10^3^, and 10^4^ PFU (n=5 per infection dose) of HPhTX NSs (**Fig 4A**) or HPhTX NSs-Nluc (**Fig 4B**) and body weight and survival were monitored daily for 14 days. Mice infected with 10^3^ and 10^4^ PFU of either virus succumbed to infection by 6– and 8-days post-infection (DPI), respectively. Mice infected with 10^2^ PFU of HPhTX NSs-Nluc exhibited 100% lethality by 10 DPI, while those infected with HPhTX NSs reached 100% lethality by 13 DPI. In mice infected with 10 PFU, 2 out of 5 mice infected with HPhTX NSs-Nluc survived infection, while only 1 out of 5 mice survived infection with HPhTX NSs. These results provide us with an estimated MLD_50_ of ∼ 15 PFU for HPhTX NSs-Nluc and a MLD_50_ of ∼ 23 PFU for HPhTX NSs that is close to the MLD_50_ of HPhTX (14). These findings suggest that HPhTX NSs and HPhTX NSs-Nluc have similar, although slightly reduced pathogenicity in C57BL/6J mice than HPhTX.

**Figure 4.**
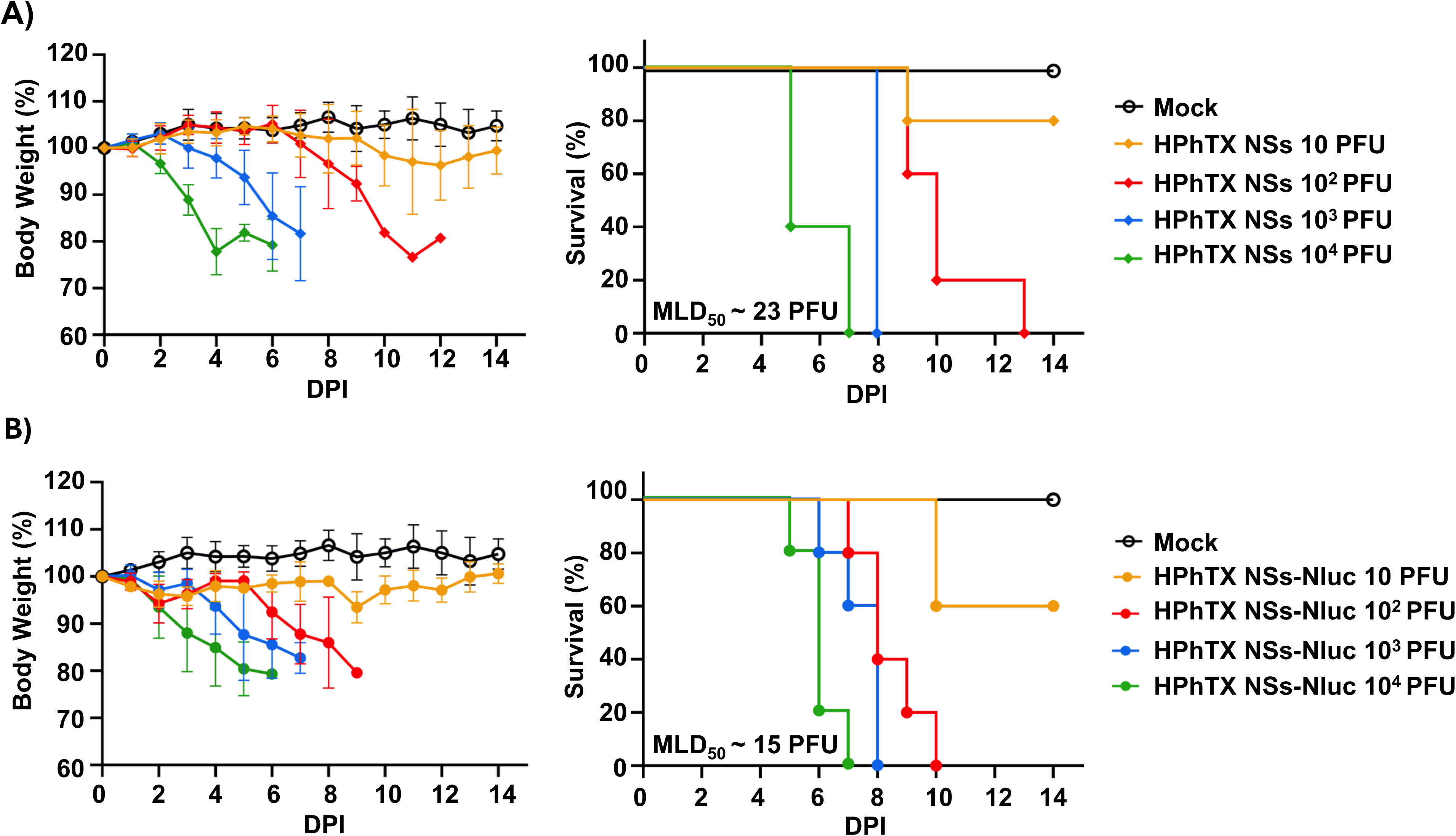
Pathogenicity of HPhTX NSs-Nluc in C57BL/6 mice. Female 6-week-old C57BL/6 mice (n=5) were inoculated with 10,10^2^,10^3^, and 10^4^ PFU of HPhTX NSs (**A**) or HPhTX NSs-Nluc (**B**) and monitored daily for 14 days for body weight (left) and survival (right). Mice that lost 25% or greater of initial weight were humanely sacrificed. The MLD_50_ was calculated by Reed and Muench method. Data represents the means and SD of the results for individual mice.

### *In vivo* tracking and *ex vivo* detection of Nluc in HPhTX NSs-Nluc-infected C57BL/6 mice

One advantage of using Nluc-expressing viruses is to track the viral infection in living animals in real-time using IVIS (9, 11, 29). This facilitates monitoring the progression of viral infection without the need to sacrifice the animals or rely on *ex vivo* imaging techniques, which can be time-consuming and require additional animal groups. To evaluate the ability to track HPhTX NSs-Nluc *in vivo*, the same animals used to evaluate viral pathogenicity were used to assess Nluc expression by IVIS on 1, 2, 4, 6, and 8 DPI (**Fig 5**). Nluc expression, which correlates with viral replication was time– and dose-dependent. As expected, the highest viral dose resulted in a stronger and earlier Nluc signal, contrary to the lower dose that had reduced signal and was detected only at 8 DPI. In mice infected with the higher doses (10^3^ and 10^4^ PFU), Nluc signal was first detected in the lungs at 2 DPI, suggesting that the virus began replicating in the respiratory system shortly after infection. By 4 DPI, Nluc signal was detected in other parts of the body, which implies that the virus spread from the lungs to other organs. We were not able to detect Nluc expression at 1 DPI, even in mice infected at the highest MOI of 10^4^ PFU. As expected, no Nluc signal was detected in the mock-infected group (**Fig 5**). To assess *ex vivo* Nluc expression, groups of 4 animals were inoculated with 10, 10^2^, 10^3^, and 10^4^ PFU of HPhTX NSs-Nluc (**Fig 6A**). Whole-body imaging confirming *in vivo* Nluc expression was performed prior to necropsy at 2 and 4 DPI (**Fig 6A**), followed by lung and brain tissue collection for *ex vivo* imaging (**Fig 6B**). In alignment with the previous data (**Fig 5**), Nluc expression was detected in the lungs as early as 2 DPI and in other organs by 4 DPI (**Fig 6A**). Likewise, Nluc expression was dose-dependent with higher levels of Nluc present in mice infected with 10^4^ PFU. Notably, we were able to detect Nluc expression *ex vivo*, in the lungs of HPhTX NSs-Nluc-infected mice at both 2 and 4 DPI. More importantly, we detected Nluc expression in the brain of some infected mice by 4 DPI, particularly those infected with the higher infection dose of 10^4^ PFU, although Nluc expression was also detected in the brain of one mouse infected with 10 and 10^2^ PFU at 4 DPI (**Fig 6B**). Surprisingly, we were not able to detect Nluc expression in the brain of any of the mice infected with 10^3^ PFU. We next quantified viral loads (**Fig 7A**) and Nluc activity (**Fig 7B**) in the NT, lungs, and brains of infected animals. Similar to the *in vivo* and *ex vivo* imaging results, we observed, a dose– and time-dependent increase in viral titers and Nluc expression (**Figs 7A and 7B, respectively**). Importantly, we observed comparable viral titers in mice infected with both viruses in all the tissues and DPI while Nluc expression was only detected in tissue samples collected from mice infected with HPhTX NSs-Nluc. These results demonstrate that HPhTX NSs-Nluc can be used for *in vivo* tracking viral infection, providing a valuable tool for studying viral dynamics in live animals with levels of Nluc expression correlating with viral titers.

**Figure 5.**
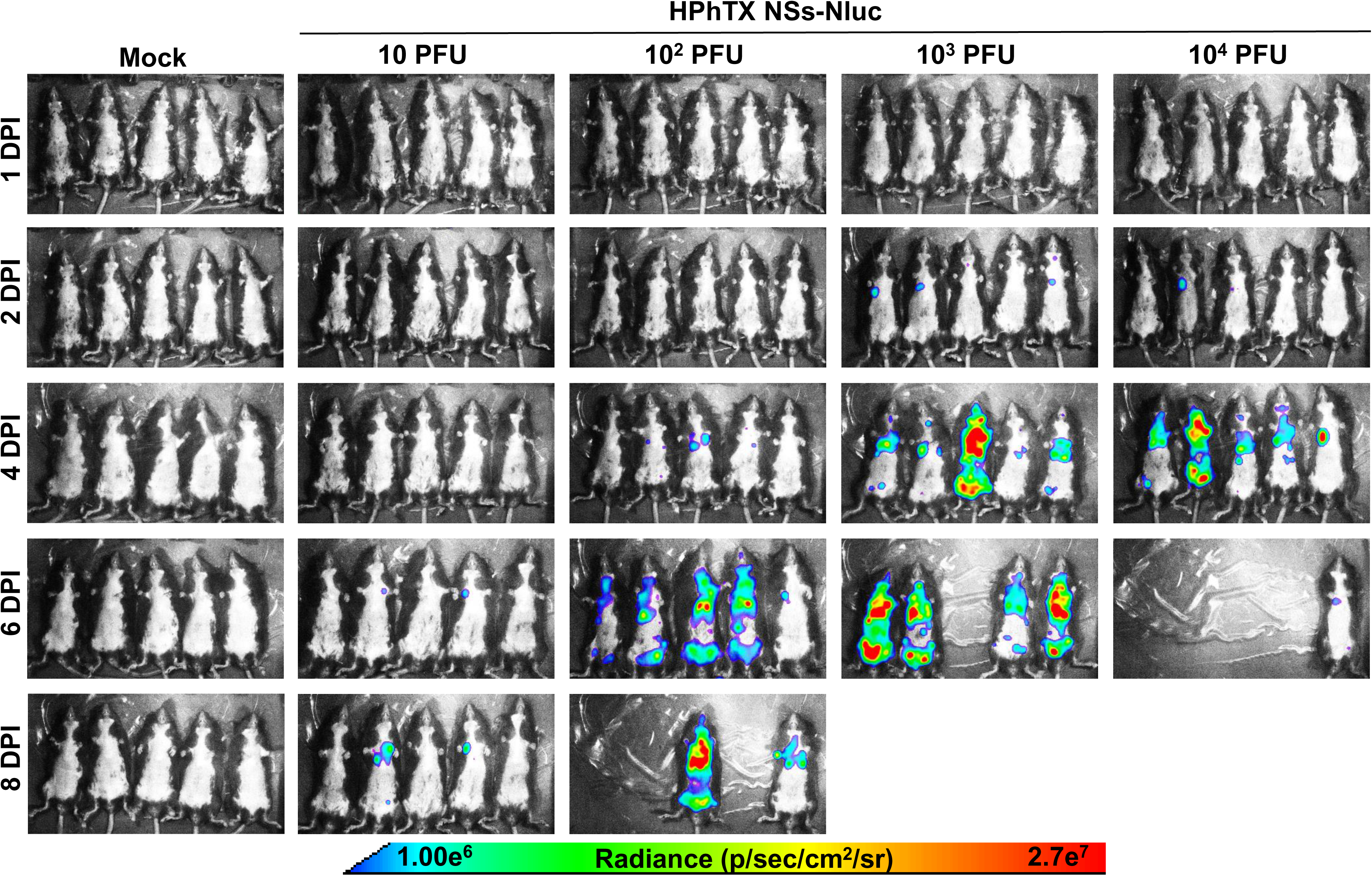
*In vivo* Nluc expression in HPhTX NSs-Nluc-infected C57BL/6 mice. Female 6-week-old C57BL/6 mice infected in Figure 4 were monitored for Nluc expression at 1, 2, 4, 6, 8 DPI using IVIS. Radiance, defined as the number of photons per s per square cm per steradian (p s−1 cm−2 sr−1), is shown on the heat map at the bottom.

**Figure 6.**
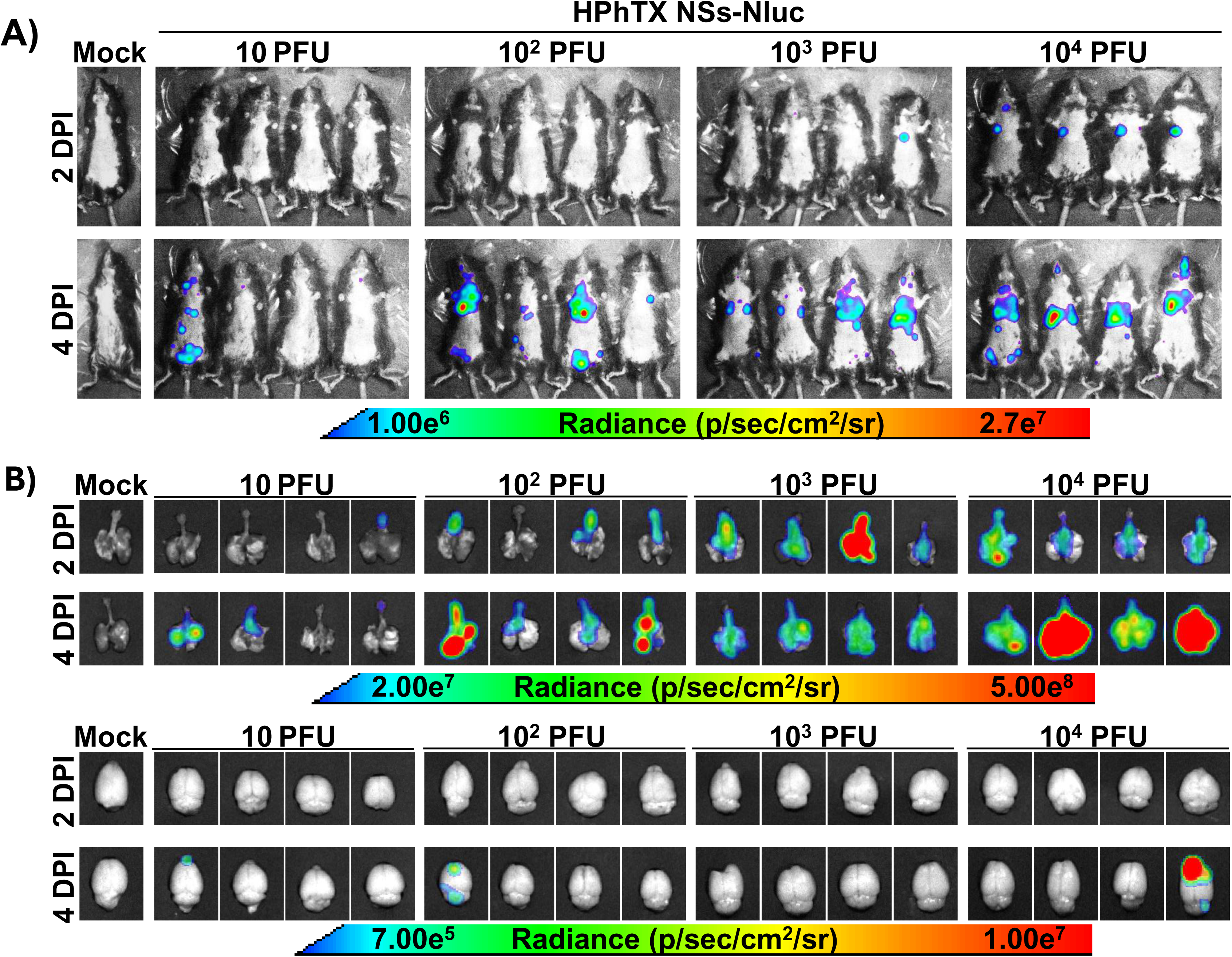
*In vivo* and *ex vivo* imaging of Nluc in C57BL/6 mice infected with HPhTX NSs-Nluc. (**A**) *In vivo* imaging of female 6-week-old C57BL/6 mice infected with the indicated doses of HPhTX NSs-Nluc at 2 and 4 DPI. (**B**) *Ex vivo* imaging of the lungs and brains of infected mice in A at 2 and 4 DPI. Radiance, defined as the number of photons per s per square cm per steradian (p s−1 cm−2 sr−1), is shown on each of the indicated heat maps.

**Figure 7.**
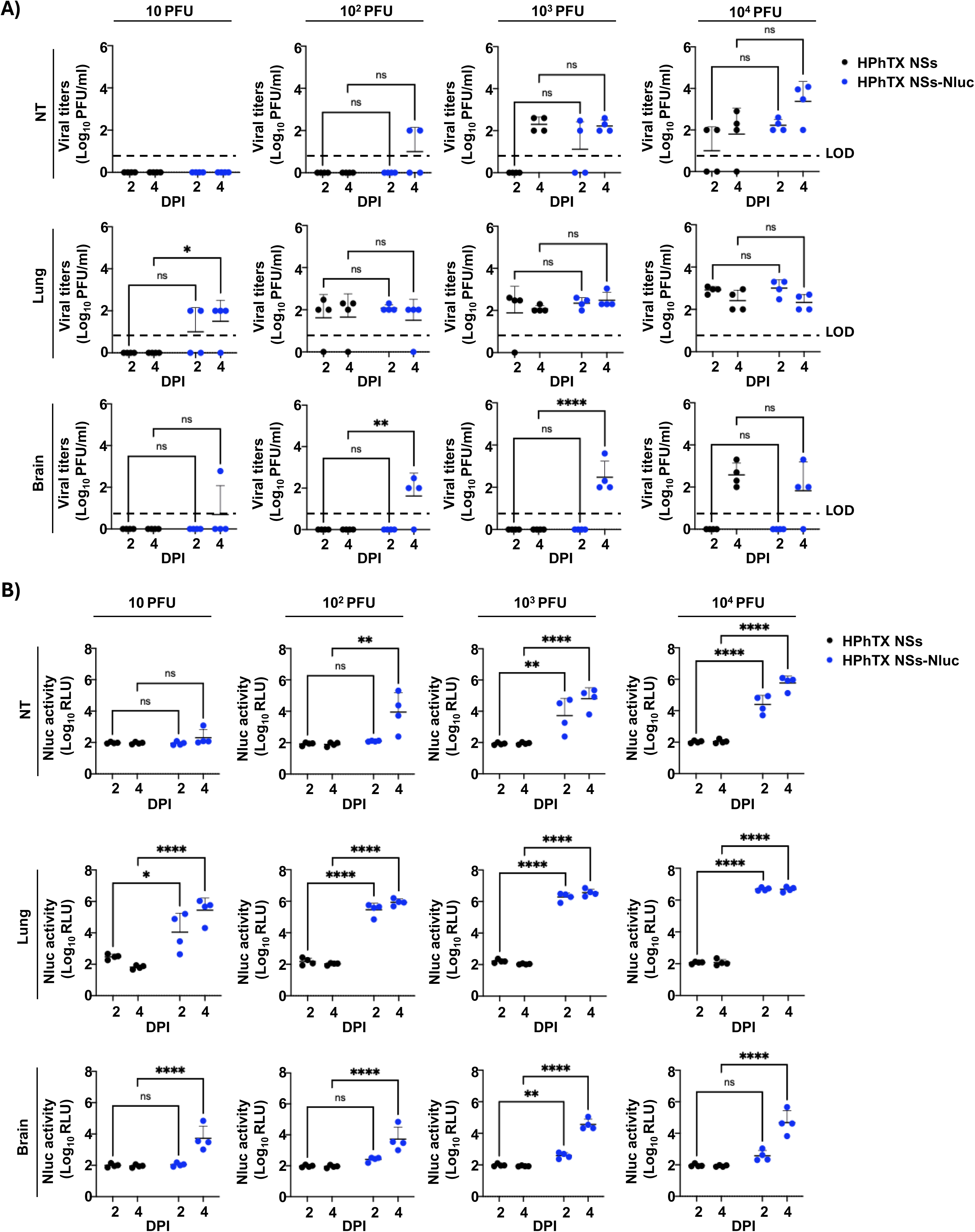
Viral titers and Nluc expression in the NT, lung and brain homogenates of HPhTX NSs-Nluc-infected C57BL/6 mice. (**A**) Viral titers in the different tissues and DPI are represented as Log 10 PFU/ml. (**B**) Nluc expression in tissue homogenates collected from the same NT, lungs, and brain tissues of mice infected with HPhTX NSs and HPhTX NSs-Nluc. The limit of detection (LOD) is indicated with a dashed line. Data are presented as mean ± SD. A two-way repeated measure ANOVA with Geisser-Greenhouse correction followed by Dunnett’s multiple comparisons test (ns=non-significant, * = *p* < 0.05, ** = *p* < 0.01, *** = *p* < 0.001, **** = *p* < 0.0001).

### *In vitro* and *in vivo* identification of therapeutics

Antiviral agents are important in protecting against viral infections, including IAVs. However, current antiviral assays to evaluate antiviral efficacy rely on secondary approaches and cannot provide real-time evaluation of antiviral activities, or longitudinal measurements. Herein, we evaluate the ability of HPhTX NSs-Nluc to identify therapeutics *in vitro* (**Fig 8**) and *in vivo* (**Figs 9-10**) using BXA, an anti-influenza drug. *In vitro*, the antiviral activity of BXA against HPhTX (IC_50_=0.016 μM), HPhTX NSs (IC_50_=0.014 μM) and HPhTX NSs-Nluc (IC_50_=0.037 μM) using a microneutralization assay was comparable (**Fig 8A**). When we determined the antiviral activity of BXA against HPhTX NSs-Nluc using a Nluc-expression assay, the IC_50_ was comparable to that observed in the microneutralization assay with HPhTX, HPhTX NSs, HPhTX NSs-Nluc (IC_50_=0.013 μM) (**Fig 8B**), demonstrating the feasibility of using Nluc expression as a valid surrogate to assess the antiviral activity against HPhTX.

**Figure 8.**
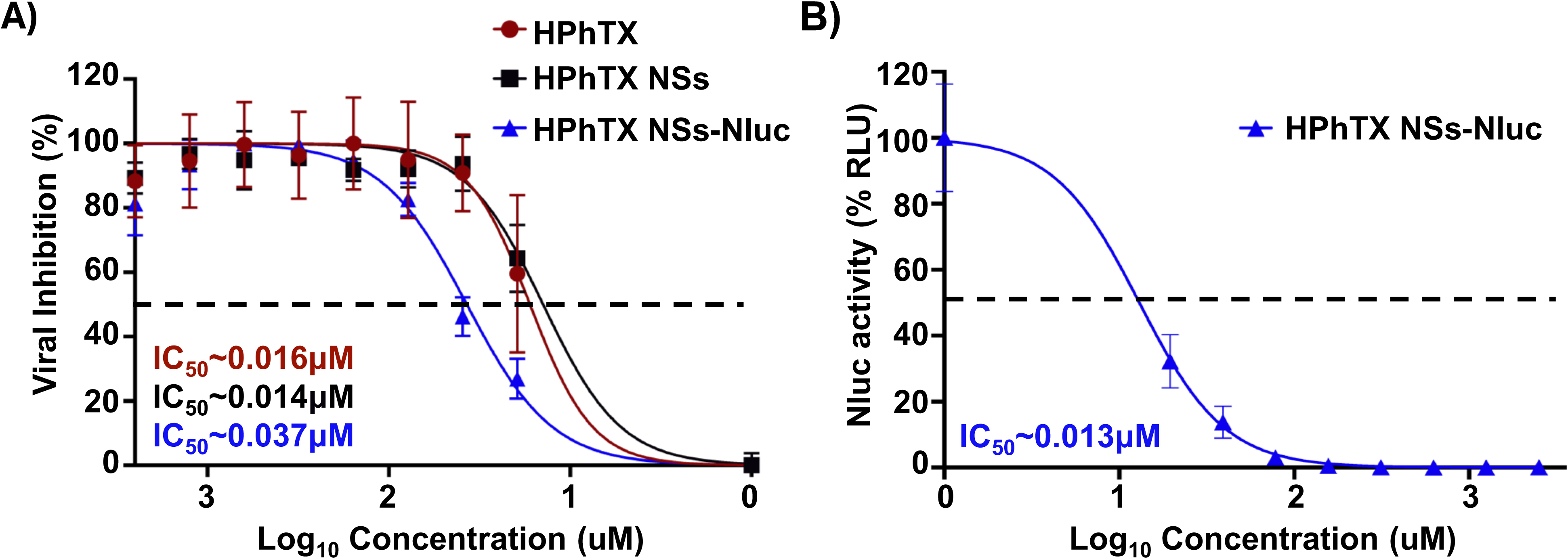
Antiviral activity of BXA against HPhTX, HPhTX NSs and HPhTX NSs-Nluc *in vitro*. The IC_50_ of BXA against HPhTX, HPhTX NSs and HPhTX NSs-Nluc was determined by microneutralization assay (**A**) or Nluc activity (**B**). The percent neutralization is calculated using sigmoidal dose-response curves. Dotted line indicates 50% of viral inhibition.

**Figure 9.**
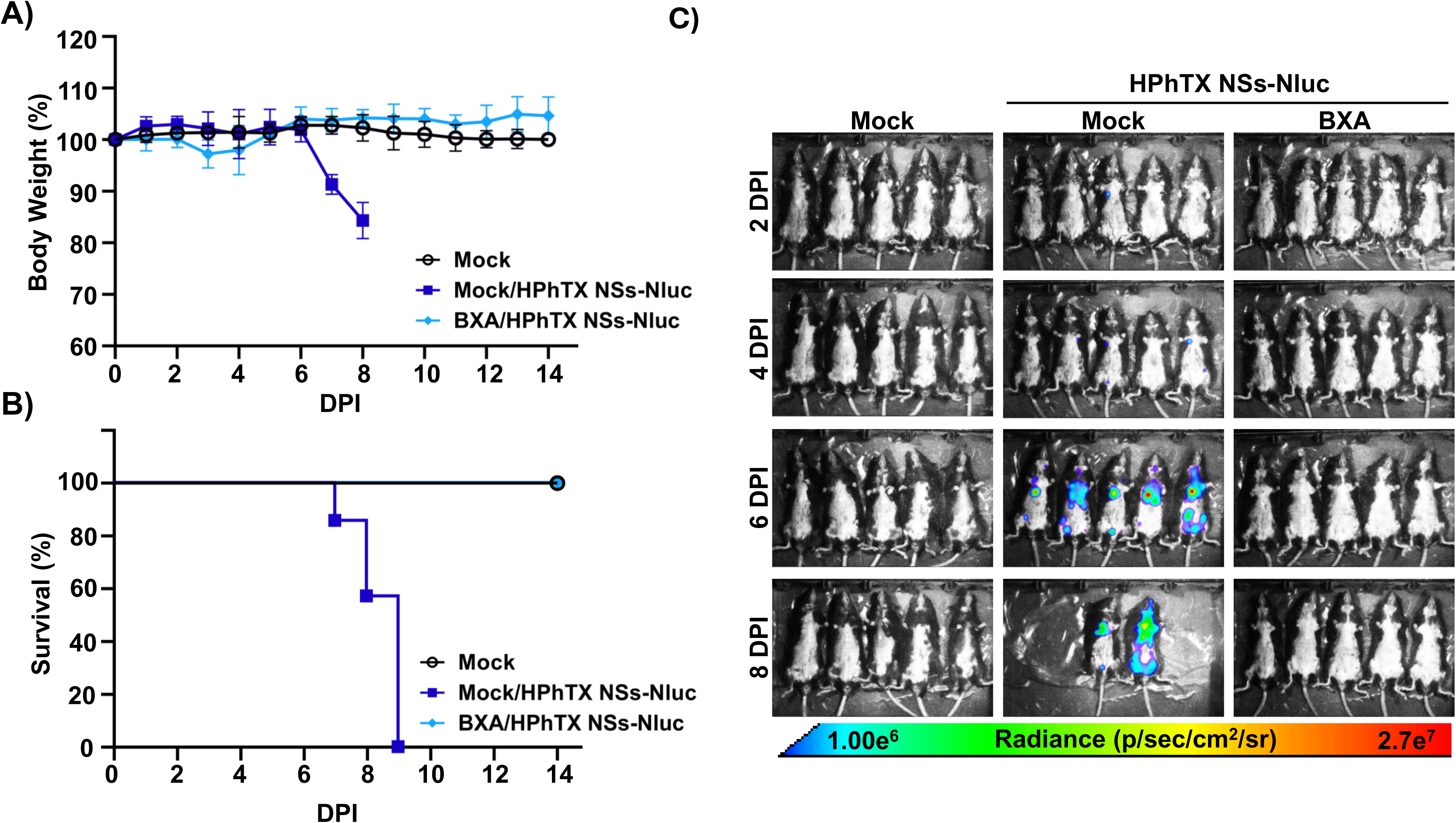
Antiviral activity of BXA against HPhTX NSs-Nluc *in vivo*. Female 6-week-old C57BL/6J mice (n=5/group) were mock-treated/mock-infected, mock-treated/infected, or treated/infected. Mice treated with BXA received 15 mg/kg of BXA twice daily by oral gavage. Mice were challenged with 10^2^ PFU of HPhTX NSs-Nluc virus at 6 h post treatment. (**A-B**) Mice were monitored for body weight changes (**A**) and survival (**B**) for 14 days. (**C**) Nluc expression in same groups of mice was monitored by IVIS on 2, 4, 6 and 8 DPI. Radiance, defined as the number of photons per s per square cm per steradian (p s−1 cm−2 sr−1), is shown on the heat map at the bottom.

**Figure 10.**
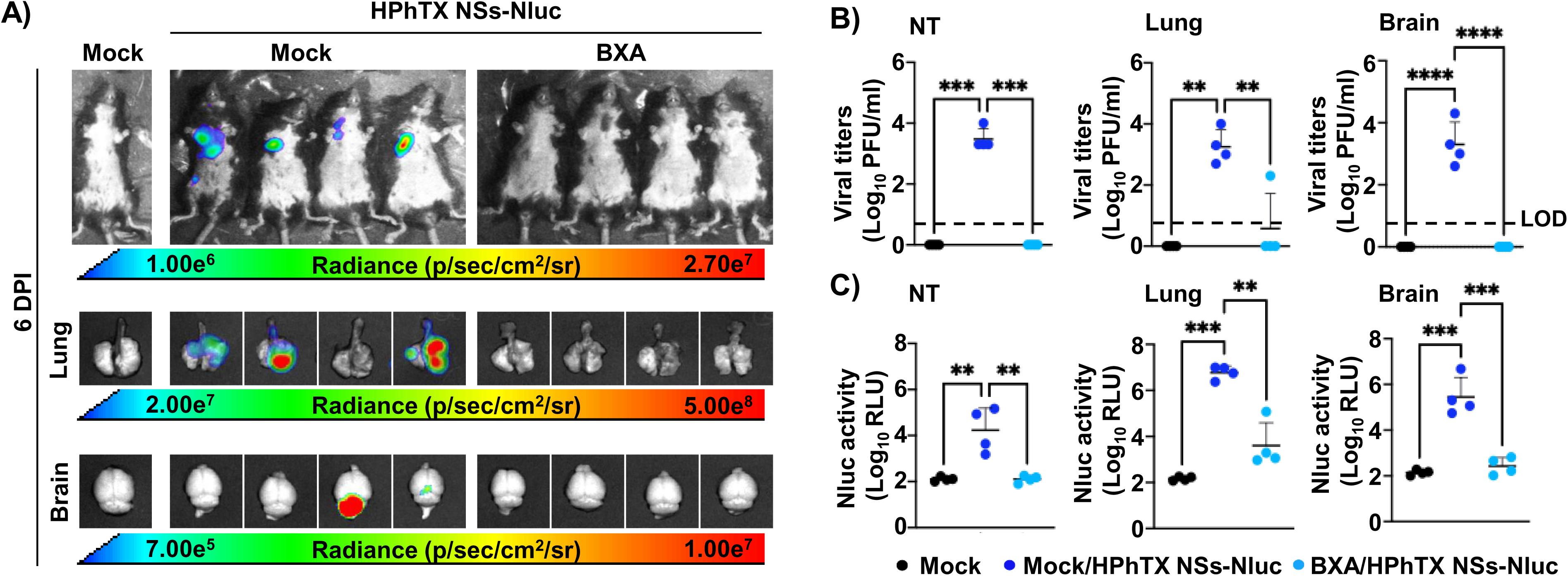
*In vivo* and *ex vivo* detection of Nluc expression and viral loads in HPhTX NSs-Nluc infected mice following treatment with BXA. (**A**) *In vivo* imaging of the entire mice and *ex vivo* imaging of lungs and brains at 6 DPI. Mice were challenged with 10^2^ PFU of HPhTX NSs-Nluc virus at 6 h post treatment. Radiance, defined as the number of photons per s per square cm per steradian (p s−1 cm−2 sr−1), is shown on the heat maps. (**B-C**) Viral titers (PFU/ml) (**B**) and Nluc expression (**C**) from the supernatants of homogenized NT, lung, and brain tissues from mice in panel A. The LOD is indicated with a dashed line. Data are presented as mean ± SD. A two-way repeated measure ANOVA with Geisser-Greenhouse correction followed by Dunnett’s multiple comparisons test (ns=non-significant, * = *p* < 0.05, ** = *p* < 0.01, *** = *p* < 0.001, **** = *p* < 0.0001).

Next, we evaluated the efficiency of using HPhTX NSs-Nluc to evaluate the antiviral activity of BXA *in vivo* (**Fig 9**). To that end, C57BL/6J mice were mock-treated or treated with BXA (15 mg/kg twice daily) and challenged i.n. with 10^2^ PFU of HPhTX NSs-Nluc. Mice treated with BXA retain their initial body weight (**Fig 9A**) with no mortalities up to 14 DPI (**Fig 9B**), and no Nluc signal at 2, 4, 6, or 8 DPI (**Fig 9C**). In contrast, all mock-treated and HPhTX NSs-Nluc-infected mice exhibited rapid body weight loss (**Fig 9A**) and all succumbed to viral infection by 9 DPI (**Fig 9B**), with high levels of Nluc expression at 6 and 8 DPI (**Fig 9C**). Viral titers and Nluc signal in the NT, lungs, and brain homogenates from another group of mice similarly treated and infected with HPhTX NSs-Nluc were determined at 6 DPI (**Fig 10**). We also evaluate Nluc signal in the same group of mice prior to necropsy (**Fig 10A**). As expected from the previous experiment, we were able to detect Nluc expression in mock-treated mice infected with HPhTX NSs-Nluc (**Fig 10A**). However, we were not able to detect Nluc expression in mice treated with BXA (**Fig 10A**). Moreover, *ex vivo* imaging of the lungs and brains (**Fig 10A**) correlated with those of *in vivo* imaging of the entire mice, with Nluc expression in the lungs of mock-treated, HPhTX NSs-Nluc-infected animals. As in our previous experiment, we were able to detect Nluc expression in the brain of two of the mock-treated animals infected with HPhTX NSs-Nluc (**Fig 10A**). However, we were not able to detect Nluc expression in the lungs or in the brains of mice treated with BXA (**Fig 10A**). Importantly, viral titers (**Fig 10B**) and Nluc expression (**Fig 10C**) in the NT, lungs and brains showed high levels of HPhTX NSs-Nluc and Nluc expression in mock-treated animals. In the case of mice treated with BXA, we only detected the presence of the virus in the lungs of one of the infected animals, with no detectable virus in the NT or in the brain (**Fig 10B**). These viral titers correlated with the presence of Nluc expression in the same tissues (**Fig 10C**). These findings demonstrate the feasibility of using HPhTX NSs-Nluc to identify therapeutics for the treatment of HPhTX infection directly *in vivo* by using IVIS.

## DISCUSSION

Reporter genes have been proven to be valuable tools for tracking viral infection *in vitro* and *in vivo*, including viral replication, tropism, and the identification of prophylactic and/or therapeutic interventions, including antivirals (9, 11–13, 30). Two main reporter genes based on fluorescence and luciferase expression have been widely used with both having advantages and disadvantages. For instance, fluorescent proteins represent an excellent option to detect the presence of the virus in infected cells using fluorescence microscopy. However, they are not optimal for live *in vivo* imaging and are restricted to *ex vivo* visualization (14). Luciferase proteins have emerged as a suitable option for live *in vivo* imaging (31). Among the different luciferase proteins, Nluc presents with several advantages, including its small size, secretory properties, ATP-independent activity, stability, and high activity, making it an ideal candidate for generating recombinant viruses (14, 34–35, 50–53).

Numerous studies, including ours, have used diverse strategies for expressing reporter genes from different viral segments (9, 13, 15). Notably, the strategy of inserting reporter genes fused to the C-terminal of the NS1 protein from a modified NS segment where the viral NS1 and NEP are separated by the PTV-1 2A autoproteolytic cleavage site has proven to be highly efficient (36–40). Therefore, we chose this approach to express Nluc from influenza A/Texas/37/2024 H5N1 (HPhTX), the first H5N1 clade 2.4.4b virus isolated from a human case in Texas. In *vitro*, HPhTX NSs-Nluc shows similar growth kinetics and plaque phenotype to that of HPhTX containing a modified NS segment (HPhTX NSs), as well as the parental HPhTX. Notably, Nluc expression was detected in all the viral plaques, confirming homogenous expression of Nluc from HPhTX NSs-Nluc. Moreover, we confirmed the stability of Nluc expression from HPhTX NSs-Nluc up to 10 serial passages in MDCK cells. Importantly, the ability of NS1 to inhibit IFNβ promoter activation when fused to Nluc was confirmed by the lack of GFP and FFluc expression in MDCK cells expressing the reporter genes under the control of an IFNβ promoter, that was comparable to that seen in cells infected with HPhTX or the HPhTX NSs encoding the modified NS segment but not in MDCK pIFNβ-GFP/IFNβ-FFluc cells infected with LPhTXdNS1, a virus lacking NS1 expression.

Our *in vivo* studies revealed that the pathogenicity of HPhTX NSs-Nluc was comparable to HPhTX NSs in C57BL/6 mice, with MLD_50_ values comparable to those from HPhTX (14), More importantly, our IVIS results demonstrate the feasibility of using HPhTX NSs-Nluc for real-time live imaging in infected C57BL/6 mice with viral titers that were dose– and time-dependent. Moreover, our *ex vivo* imaging results demonstrate the feasibility of using HPhTX NSs-Nluc to detect the presence of the virus in the lungs and brains of infected mice. However, while we were able to detect the presence of the virus, as determined by Nluc expression, in the lungs of all infected mice, we were only able to detect Nluc expression in the brain of some infected animals. Neurotropism is a unique characteristic of HPAIV H5N1 viruses. We are currently trying to determine the limitations of detecting Nluc expression in the brain of infected mice and the potential reasons for the lack of reproducibility in our initial studies. Importantly, quantification of viral titers and Nluc expression in tissue homogenates from lungs and brains correlate with levels of Nluc expression *in vivo* and *ex vivo*, including similar dose– and time-dependent increase, consistent with HPhTX NSs, or HPhTX infection (14).

Another advantage of viruses expressing reporter genes is their use to interrogate libraries of compounds and/or biologicals to identify those with antiviral and/or neutralizing activity in high-throughput screening (HTS) settings. Here we demonstrate the feasibility of using HPhTX NSs-Nluc to identify antivirals *in vitro* by assessing Nluc expression that can be implemented for the screening of large libraries to identify those with antiviral activity. More importantly, we also demonstrate the ability of HPhTX NSs-Nluc to identify therapeutics *in vivo* where Nluc expression can be used as a valid surrogate to determine the ability of a compound to inhibit viral infection. Moreover, since Nluc can be tracked from the same animals at different times post-infection, this reduces the number of mice needed to identify antivirals, opening the possibility for *in vivo* screening of antiviral drugs.

## Authors Contributions

Conceptualization: A.M. and L.M-S.; Methodology: R.S.B., A.M., R.E., E.C., R.N.P., A.C., M.B., N.J. and C.Y.; Data collection and interpretation: A.M., R.S.B., T.J.C.A., A.G-S., and L.M-S.; Funding acquisition and resources: A.M., T.J.C.A., A.G-S., and L.M-S.; Writing— original draft preparation: R.S.B., A.M. and L.M-S.; Writing—review and editing: all authors have read and agreed to the published version of the manuscript.

## References

1. Mostafa A, Abdelwhab EM, Mettenleiter TC, Pleschka S. 2018. Zoonotic Potential of Influenza A Viruses: A Comprehensive Overview. Viruses 10.

2. Youk S, Torchetti MK, Lantz K, Lenoch JB, Killian ML, Leyson C, Bevins SN, Dilione K, Ip HS, Stallknecht DE, Poulson RL, Suarez DL, Swayne DE, Pantin-Jackwood MJ. 2023. H5N1 highly pathogenic avian influenza clade 2.3.4.4b in wild and domestic birds: Introductions into the United States and reassortments, December 2021–April 2022. Virology 587:109860.

3. Mostafa A, Naguib Mahmoud M, Nogales A, Barre Ramya S, Stewart James P, García-Sastre A, Martinez-Sobrido L. 2024. Avian influenza A (H5N1) virus in dairy cattle: origin, evolution, and cross-species transmission. mBio 15:e02542–24.

4. Halwe NJ, Cool K, Breithaupt A, Schön J, Trujillo JD, Nooruzzaman M, Kwon T, Ahrens AK, Britzke T, McDowell CD, Piesche R, Singh G, Pinho dos Reis V, Kafle S, Pohlmann A, Gaudreault NN, Corleis B, Ferreyra FM, Carossino M, Balasuriya UBR, Hensley L, Morozov I, Covaleda LM, Diel DG, Ulrich L, Hoffmann D, Beer M, Richt JA. 2025. H5N1 clade 2.3.4.4b dynamics in experimentally infected calves and cows. Nature 637:903–912.

5. CDC. 2025. H5 Bird Flu: Current Situation. https://www.cdc.gov/bird-flu/situation-summary/index.html. Accessed 02.18.2025.

6. USDA. 2025. The Occurrence of Another Highly Pathogenic Avian Influenza (HPAI) Spillover from Wild Birds into Dairy Cattle.

7. Nogales A, Schotsaert M, Rathnasinghe R, DeDiego ML, García-Sastre A, Martinez-Sobrido L. 2021. Replication-Competent ΔNS1 Influenza A Viruses Expressing Reporter Genes. Viruses 13.

8. Martinez-Sobrido L, Nogales A. 2024. Recombinant Influenza A Viruses Expressing Reporter Genes from the Viral NS Segment. Int J Mol Sci 25.

9. Nogales A, Ávila-Pérez G, Rangel-Moreno J, Chiem K, DeDiego ML, Martínez-Sobrido L. 2019. A Novel Fluorescent and Bioluminescent Bireporter Influenza A Virus To Evaluate Viral Infections. J Virol 93.

10. Breen M, Nogales A, Baker SF, Perez DR, Martínez-Sobrido L. 2016. Replication-Competent Influenza A and B Viruses Expressing a Fluorescent Dynamic Timer Protein for In Vitro and In Vivo Studies. PLoS One 11:e0147723.

11. Chiem K, Rangel-Moreno J, Nogales A, Martinez-Sobrido L. 2019. A Luciferase-fluorescent Reporter Influenza Virus for Live Imaging and Quantification of Viral Infection. J Vis Exp doi:10.3791/59890.

12. Breen M, Nogales A, Baker SF, Martínez-Sobrido L. 2016. Replication-Competent Influenza A Viruses Expressing Reporter Genes. Viruses 8.

13. Nogales A, Rodríguez-Sánchez I, Monte K, Lenschow DJ, Perez DR, Martínez-Sobrido L. 2016. Replication-competent fluorescent-expressing influenza B virus. Virus Res 213:69–81.

14. Mostafa A, Barre RS, Allué-Guardia A, Escobedo RA, Shivanna V, Rothan H, Castro EM, Ma Y, Cupic A, Jackson N, Bayoumi M, Torrelles JB, Ye C, García-Sastre A, Martinez-Sobrido L. 2025. Replication kinetics, pathogenicity and virus-induced cellular responses of cattle-origin influenza A(H5N1) isolates from Texas, United States. Emerging Microbes & Infections 14:2447614.

15. Nogales A, Baker SF, Martínez-Sobrido L. 2015. Replication-competent influenza A viruses expressing a red fluorescent protein. Virology 476:206–216.

16. Martínez-Sobrido L, García-Sastre A. 2010. Generation of recombinant influenza virus from plasmid DNA. J Vis Exp doi:10.3791/2057.

17. Mostafa A, Kanrai P, Ziebuhr J, Pleschka S. 2013. Improved dual promotor-driven reverse genetics system for influenza viruses. J Virol Methods 193:603–10.

18. Mostafa A, Kanrai P, Petersen H, Ibrahim S, Rautenschlein S, Pleschka S. 2015. Efficient Generation of Recombinant Influenza A Viruses Employing a New Approach to Overcome the Genetic Instability of HA Segments. PLOS ONE 10:e0116917.

19. Mostafa A, Kanrai P, Ziebuhr J, Pleschka S. 2016. The PB1 segment of an influenza A virus H1N1 2009pdm isolate enhances the replication efficiency of specific influenza vaccine strains in cell culture and embryonated eggs. J Gen Virol 97:620–631.

20. Quinlivan M, Zamarin D, García-Sastre A, Cullinane A, Chambers T, Palese P. 2005. Attenuation of equine influenza viruses through truncations of the NS1 protein. J Virol 79:8431–9.

21. O’Neill RE, Talon J, Palese P. 1998. The influenza virus NEP (NS2 protein) mediates the nuclear export of viral ribonucleoproteins. Embo j 17:288–96.

22. Steinig E, Coin LJJoOSS. 2022. Nanoq: ultra-fast quality control for nanopore reads. 7:2991.

23. Li H. 2016. Minimap and miniasm: fast mapping and de novo assembly for noisy long sequences. Bioinformatics 32:2103–10.

24. Li H, Handsaker B, Wysoker A, Fennell T, Ruan J, Homer N, Marth G, Abecasis G, Durbin R. 2009. The Sequence Alignment/Map format and SAMtools. Bioinformatics 25:2078–9.

25. Pedersen BS, Quinlan AR. 2018. Mosdepth: quick coverage calculation for genomes and exomes. Bioinformatics 34:867–868.

26. Wilm A, Aw PP, Bertrand D, Yeo GH, Ong SH, Wong CH, Khor CC, Petric R, Hibberd ML, Nagarajan N. 2012. LoFreq: a sequence-quality aware, ultra-sensitive variant caller for uncovering cell-population heterogeneity from high-throughput sequencing datasets. Nucleic Acids Res 40:11189–201.

27. Hegazy A, Soltane R, Alasiri A, Mostafa I, Metwaly AM, Eissa IH, Mahmoud SH, Allayeh AK, Shama NMA, Khalil AA, Barre RS, El-Shazly AM, Ali MA, Martinez-Sobrido L, Mostafa A. 2024. Anti-rheumatic colchicine phytochemical exhibits potent antiviral activities against avian and seasonal Influenza A viruses (IAVs) via targeting different stages of IAV replication cycle. BMC Complementary Medicine and Therapies 24:49.

28. Khalil AM, Nogales A, Martínez-Sobrido L, Mostafa A. 2024. Antiviral responses versus virus-induced cellular shutoff: a game of thrones between influenza A virus NS1 and SARS-CoV-2 Nsp1. Frontiers in Cellular and Infection Microbiology 14.

29. Smith S, Rayner JO, Kim JH. 2025. Fluorofurimazine, a novel NanoLuc substrate, enhances real-time tracking of influenza A virus infection without altering pathogenicity in mice. Microbiol Spectr 13:e0268924.

30. Kim JH, Bryant H, Fiedler E, Cao T, Rayner JO. 2022. Real-time tracking of bioluminescent influenza A virus infection in mice. Scientific Reports 12:3152.

31. Liu S, Su Y, Lin MZ, Ronald JA. 2021. Brightening up Biology: Advances in Luciferase Systems for in Vivo Imaging. ACS Chem Biol 16:2707–2718.

